# Conspecific Presence Facilitates the Reliable Expression of Nicotine Reward in Juvenile Zebrafish

**DOI:** 10.64898/2026.06.17.732931

**Authors:** Jun Huang, Thirumalini Vaithianathan, Hao Chen

## Abstract

**Rationale:** Adolescence is a period of heightened vulnerability to nicotine reinforcement. While zebrafish are a valuable model for investigating drug reward, standard conditioned place preference (CPP) assays typically test subjects in isolation. In this highly social species, solitary testing may act as an environmental stressor that confounds behavioral readouts.

**Objectives:** This study examined how social context during testing (isolated vs. grouped) affects experimental attrition, behavioral stability, and nicotine CPP expression in late juvenile zebrafish.

**Methods:** Zebrafish housed in groups of four were tested either individually (isolated) or in their housing groups (grouped) during daily 20-minute sessions. Following baseline preference assessments, subjects underwent six days of conditioning pairing their initially non-preferred compartment with fish water or nicotine (0.5, 1.6, or 5.0 µmol/L). Place preference, locomotion, and thigmotaxis were assessed on a drug-free test day.

**Results:** Isolated testing reduced distance traveled, decreased swimming speed, and increased time spent near tank walls, indicating heightened anxiety-like behavior. Experimental attrition was significantly higher in isolated (38.9%) than grouped (2.5%) subjects. Grouped subjects developed significant place preference at 1.6 and 5.0 µ mol/L nicotine, whereas preference was not detectable in isolated subjects.

**Conclusions:** Solitary testing acts as a stressor that increases experimental attrition and masks place preference. Conversely, testing in the presence of conspecifics stabilizes behavior and facilitates the detection of nicotine reward in late juvenile zebrafish.

## Introduction

Adolescence is a developmental period of heightened vulnerability to the reinforcing effects of addictive substances. This susceptibility is attributed, in part, to asynchronous maturation of brain circuits: subcortical reward networks, including the mesolimbic dopamine system, mature early, whereas prefrontal regions responsible for inhibitory control develop more slowly (Counotte et al. 2011; Leslie 2020). In rodent models, adolescent animals acquire drug self-administration across a broader range of doses than adults and show reduced sensitivity to the aversive, intake-limiting properties of nicotine and psychostimulants (Chen et al. 2006; Natividad et al. 2013). Whether analogous developmental sensitivities exist in non-mammalian vertebrates remains an active area of investigation.

Zebrafish (*Danio rerio*) have become a widely used model organism in neurobehavioral pharmacology because of their conserved monoaminergic neurotransmitter systems and high genetic homology with mammals. Conditioned place preference (CPP) is the principal behavioral assay for measuring the reinforcing properties of drugs in zebrafish (Kedikian et al. 2013; Viscarra et al. 2020). Although CPP has been validated extensively in adult zebrafish (>90 days of age), its application to younger, juvenile stages remains challenging. Behavioral assays in juvenile zebrafish are constrained by high locomotor variability, elevated experimental attrition, and a narrow effective concentration window for nicotine: rewarding effects have been reported within the 0.63 to 6.3 µmol/L range, whereas higher concentrations elicit avoidance or sensory irritation (Schneider et al. 2023). Nicotine is of particular interest for studying social-environmental interactions because its stimulus profile comprises both reinforcing and aversive components. In adolescent rodents, peer social interactions facilitate the acquisition of nicotine self-administration by helping subjects overcome the drug’s initial aversive sensory properties (Chen et al. 2011). Whether social context exerts a comparable influence on behavioral responses to nicotine in juvenile zebrafish has not been investigated.

The social environment is a critical, yet often uncontrolled, variable in behavioral pharmacology assays with juvenile zebrafish. Zebrafish are obligate shoaling fish that rely on group interactions for defense, foraging, and predator avoidance. Social isolation is a well-characterized stressor in this species, although sensitivity to isolation varies across development (Petersen et al. 2022). In mammals, the presence of conspecifics can attenuate the behavioral and physiological consequences of stress, often referred to as social buffering (Kikusui et al. 2006); this process is less well characterized in teleost fish. Larval zebrafish (6 days post-fertilization) show minimal behavioral or endocrine responses to brief separation from conspecifics (Bai et al. 2016). In contrast, early juveniles (2 to 3 weeks of age) are highly sensitive to social isolation, which manifests as reduced locomotion, impaired social preference, and frequent freezing (Tunbak et al. 2020). By the late juvenile stage (approximately 60 to 65 days of age), shoaling behavior is fully established, yet sexual maturation and neural development are still ongoing(Parichy et al. 2009; Singleman and Holtzman 2014). This developmental window may therefore be especially sensitive to the stress of social isolation during behavioral testing. In adult zebrafish (>90 days of age), the behavioral effects of short-term isolation are more variable, and endocrine stress responses can decouple from classical anxiety-like behaviors (Marchetto et al. 2021; Onarheim et al. 2022).

The present study investigated how social context affects the feasibility and expression of nicotine-induced CPP in juvenile AB strain zebrafish by comparing subjects tested in groups of four with subjects tested individually. We predicted that the presence of conspecifics would reduce isolation stress, stabilize exploratory behavior, and facilitate CPP expression, whereas solitary testing would increase anxiety-like behavior and mask conditioned place preference.

## Methods

### Animals

Late juvenile AB strain zebrafish (Danio rerio, approximately 60 to 65 days of age at the beginning of testing) were used in this study. The animals were ordered as embryos from the Zebrafish International Resource Center (University of Oregon) and reared in-house. All subjects were raised in tanks containing 10 to 20 individuals. A total of 117 subjects (with approximately similar numbers of males and females) were used. Subjects were housed in groups of 4 during the experiment and were tested either in groups of four or individually. To permit individual identification and longitudinal tracking, subjects tested in isolation were tagged using visible implant elastomer (Hohn and Petrie-Hanson 2013) at least three days before the first baseline session. Subjects were returned to their housing tanks immediately after each session. All fish were maintained under standard husbandry conditions prior to testing.

### Apparatus

The CPP tanks (226 mm × 90 mm × 113 mm) were 3D printed using white Polylactic acid filament. The chamber floor was constructed with the opaque plexiglass and divided into two visually distinct zones defined by different floor cues: three large black dots on the left side and a grid pattern on the right side. Illumination from below was provided by two LED panels controlled by a Raspberry Pi computer. Behavior was recorded at 30 frames/s using a Raspberry Pi camera mounted above the test chamber. Detailed chamber specifications, including files for the floor patterns, are available at https://github.com/miraclezero/FishCPP. For each testing session, each tank was filled with 1 L of system water. Multiple tanks were tested simultaneously, and visual barriers prevented fish from seeing individuals in adjacent tanks.

### Experimental Protocol

A biased CPP protocol was used to evaluate the reinforcing effects of nicotine. The experimental design consisted of a two-day baseline period, a six-day conditioning phase, and a final preference test. All baseline and conditioning sessions were conducted in the morning, approximately at 11 AM. Conditioning testing was conducted at approximately 4 PM.

During the baseline period, subjects had free access to both compartments of the chamber, one with three black dots and one with a black grid floor cue. For each individual (isolated condition) or tank (grouped condition), the compartment in which the subjects spent less than 50% of their time during Baseline 2 was designated as the initially non-preferred compartment, while the preferred compartment was designated as the unpaired compartment. Because there was no strong population-level innate bias for either cue, the dots and grid patterns were counterbalanced as the paired and unpaired stimuli across subjects.

Subjects then underwent six daily conditioning sessions, where a physical divider was inserted to restrict access to one compartment at a time. The session consisted of two sequential 20-minute phases. Fish were initially confined to their initially preferred compartment for 20 minutes in fish water. Immediately following the first phase, fish were confined to their initially non-preferred compartment for 20 minutes in the presence of either nicotine (0.5, 1.6, or 5.0 µmol/L bath concentration) or fish water (control group). These nicotine concentrations were selected based on prior behavioral evidence showing that rewarding effects in young zebrafish occur within a narrow, low-micromolar window (0.63 to 6.3 µmol/L), whereas higher concentrations trigger avoidance or physical distress (Schneider et al. 2023).

A final CPP test was then conducted in the afternoon of the last conditioning session. The physical divider was removed, allowing subjects free access to both compartments of the chamber for 20 minutes in the absence of nicotine to measure the post-conditioning preference shift.

### Video Acquisition and Tracking

Each experimental session was recorded for 20 minutes from an overhead perspective using Raspberry Pi computers equipped with Pi Camera modules at a frame rate of 30 frames/s. To allow the subjects to habituate to the chamber, the first 5 minutes of each video recording were skipped from analysis, resulting in a 15-minute analysis window. The resulting video files were processed using the SLEAP (Social LeAP) deep learning software (version 1.4.1a2) (Pereira et al. 2022). For each frame, the tracking model identified and extracted the spatial coordinates of each subject. For grouped cohorts, the model detected all four subjects simultaneously. Although individual identity was not tracked across daily sessions, coordinate trajectories within each session were checked for identity flips (tracking swaps between subjects) to maintain accurate coordinate histories for all individuals. These pixel coordinates were then converted to physical distances using a conversion factor of 0.149 mm/pixel. To filter out high-frequency coordinate noise (tracking jitter) and prevent the accumulation of artificial displacement when the subjects were stationary (Noldus et al. 2001), trajectories were smoothed using a centered moving average filter with a rolling window of 30 frames (1.0 second), and a minimum frame-to-frame displacement threshold of 0.5 mm was applied. The filtered trajectories were subsequently analyzed to calculate the total distance traveled, mean swimming speed, compartment occupancy, and thigmotaxis during each session. Thigmotaxis was defined as the percentage of time spent in the peripheral zone within 10 mm of the tank walls.

### Data Analysis and Statistics

To quantify the shift in preference, a place preference score was calculated as the difference between the proportion of time spent in the paired compartment during the post-conditioning test and that spent during the baseline session. For isolated subjects, this score was calculated at the individual level using the second baseline session (B2) as the baseline reference. For grouped subjects, because individual identity could not be tracked across independent daily sessions, the place preference score was calculated using the average baseline 2 preference of the group (tank-mean preference) as the reference. This group-mean baseline reference was justified by the high behavioral cohesion and low variance observed between individual subjects within each tank.

Behavioral cohesion among grouped subjects was characterized at two levels. At the tank level, social cohesion was quantified as the standard deviation of the preference ratio (*P*) among the four fish within each tank; a lower value indicates that tankmates adopted more similar spatial preferences during a given session. At the population level, group cohesion was estimated using the Intraclass Correlation Coefficient (ICC) derived from the LMM random intercept for Tank ID; the ICC represents the proportion of total behavioral variance attributable to tank membership across all grouped subjects. The overall population-level dispersion was compared between testing conditions using Levene’s test and F-test. Behavioral stability was assessed using Pearson correlation between the two baseline sessions (B1 and B2). For grouped subjects, because individual identity was not tracked across daily sessions, preference ratios were averaged across all four fish in each tank to obtain tank-level group means for each session, and the correlation was calculated on these tank averages. For isolated subjects, since individuals were tested alone and their identities were known, the correlation was calculated directly on individual session preferences.

To account for shared environmental variance in grouped subjects (potential correlation of behavior among the four fish in a tank) and differences in variability between testing conditions, longitudinal locomotor metrics (total distance traveled and mean swimming speed) were analyzed using a Linear Mixed-Effects Model (LMM) with Tank ID as a random intercept and heterogeneous residual variances modeled across testing conditions (represented by the factor *Housing* in the statistical model, which encodes the social context during testing: isolated vs. grouped):

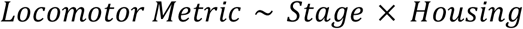

with random intercept (1|*Tank ID*) and residual variance weighting by *Housing*. This heterogeneous LMM was fitted using the nlme package in R, which supports explicit modeling of unequal residual variances across groups; the lme4 package does not provide this functionality. For CPP scores, which were analyzed cross-sectionally and did not require heterogeneous variance structures, standard Linear Models (LM) were utilized for isolated subjects, and random-intercept LMM were utilized for grouped subjects to account for shared tank environment:

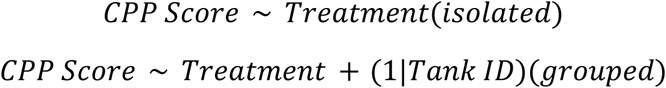

All CPP models were fitted in R using the lme4 and lmerTest packages, with degrees of freedom and p-values computed using Satterthwaite’s approximation. Post-hoc comparisons between specific treatment doses and controls were performed using Dunnett’s adjustment. To verify that LMM random intercepts did not inflate statistical precision, a sensitivity analysis was also conducted by collapsing individual scores to tank averages and fitting a standard linear model on the tank means (*N* = 20 tanks). As a secondary analysis, we tested whether each treatment group’s CPP score differed significantly from zero (i.e., whether the group shifted from its own baseline). For grouped subjects, this was assessed by fitting an intercept-only LMM per treatment (*CPP Score* ∼ 1 + (1|*Tank ID*)) and testing the intercept. For isolated subjects, one-sample t-tests were used.

## Results

### Sample Sizes and Experimental Attrition

A total of 117 juvenile zebrafish were initially tested, with 81 subjects studied in groups of four and 36 subjects evaluated individually. Of these, 101 subjects completed the final place preference test, while 16 subjects were excluded due to immobility or escape behaviors. The attrition rate was much higher in the isolated condition (38.9%, 14 out of 36 fish excluded) than in the grouped condition (2.5%, 2 out of 81 fish excluded). Table 1 outlines the initial, excluded, and completed sample sizes, as well as the attrition rates, across all testing conditions and treatment groups.

**Table 1.** Sample size and experimental attrition across testing conditions and treatment groups.

| Testing Condition | Treatment Group | Initial (N) | Excluded (N) | Completed (N) | Attrition Rate (%) |
| --- | --- | --- | --- | --- | --- |
| **Grouped** (4 fish/tank) | Control | 28 | 0 | 28 | 0.0% |
|  | 0.5 µmol/L Nicotine | 20 | 1 | 19 | 5.0% |
|  | 1.6 µmol/L Nicotine | 20 | 0 | 20 | 0.0% |
|  | 5.0 µmol/L Nicotine | 13 | 1 | 12 | 7.7% |
|  | *Grouped Total* | *81* | *2* | *79* | *2.5%* |
| **Isolated** (1 fish/tank) | Control | 16 | 6 | 10 | 37.5% |
|  | 1.6 µmol/L Nicotine | 20 | 8 | 12 | 40.0% |
|  | *Isolated Total* | *36* | *14* | *22* | *38.9%* |

### Baseline Preference, Behavioral Stability, and Social Cohesion

Initial preference and spatial exploration were characterized during the two baseline sessions (B1 and B2). Track density plots revealed that subjects tested in isolation exhibited marked thigmotaxis, remaining near the perimeter of the tank and showing restricted spatial exploration. In contrast, grouped subjects explored both zones of the chamber uniformly, displaying a balanced spatial distribution across the arena (**Figure 3**).

**Figure 1.**
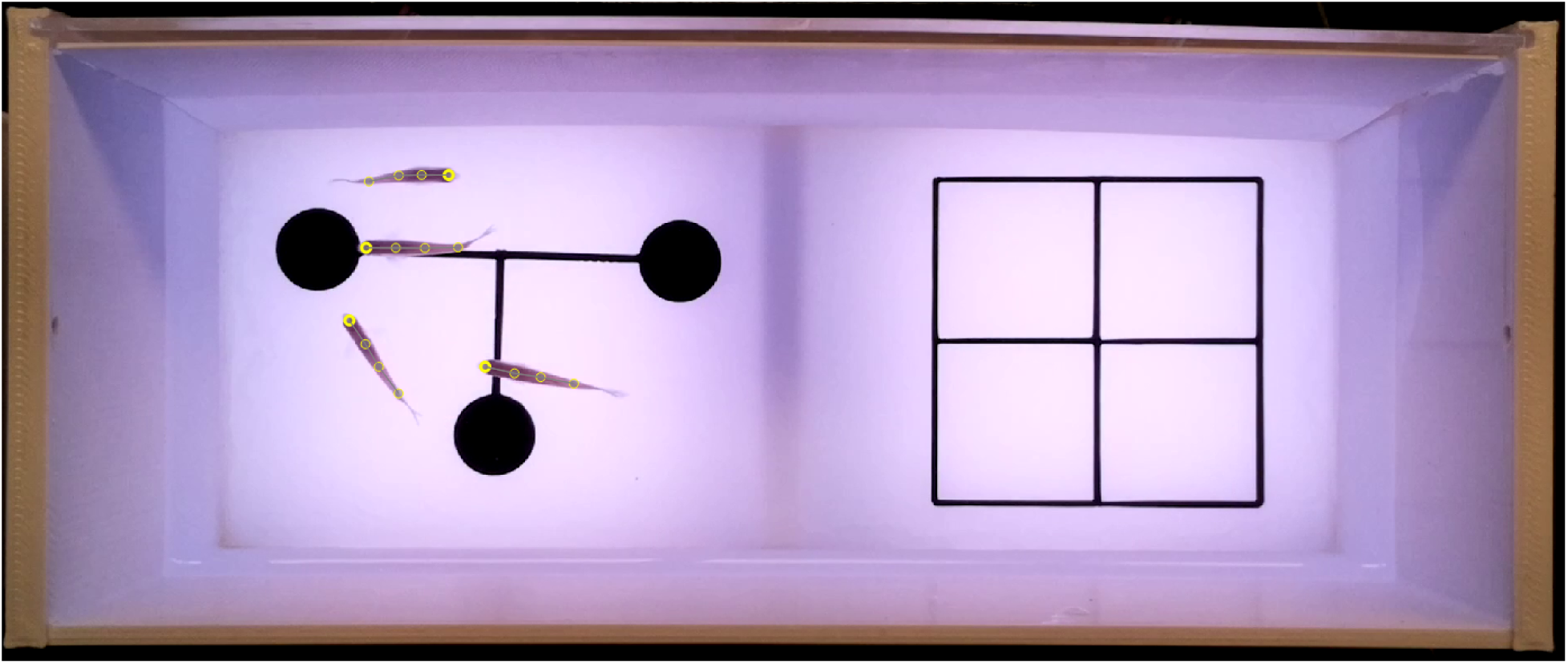
Experimental arena setup. The tank features two zones with distinct visual cues (dots and grids) to measure place preference.

**Figure 2.**
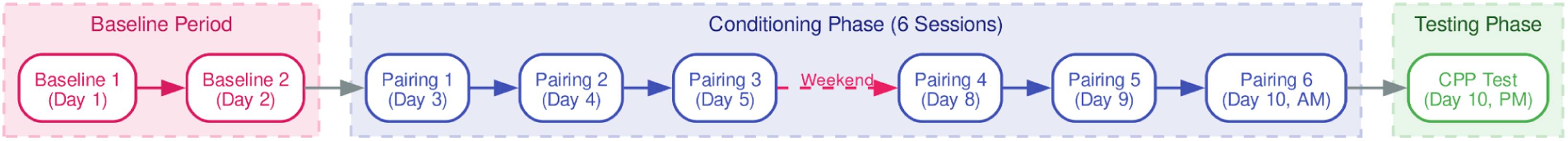
Experimental Protocol. Timeline showing the sequence of baseline tests, conditioning sessions, and the final preference test.

**Figure 3.**
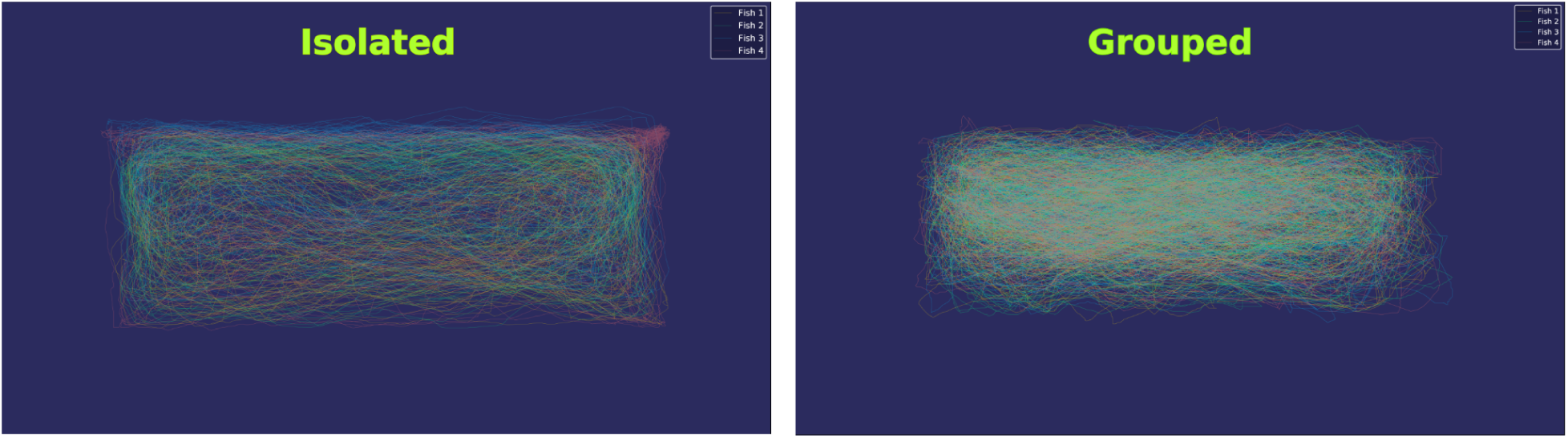
Locomotor trajectories in the place preference chamber. Overhead track density plots at Baseline 2 show typical spatial exploration for four fish tested individually with their tracks combined (left) and four fish tested together as a group (right). Isolated subjects exhibited thigmotaxis (a tendency to remain near the perimeter of the tank) and restricted exploration, whereas grouped subjects explored both zones uniformly.

### Thigmotaxis and Anxiety-like Behavior

To evaluate the impact of the social context during testing on anxiety-like behavior, we analyzed thigmotaxis levels across experimental stages and treatment groups. LMM analysis revealed a pervasive effect of the testing condition (social context) on thigmotaxis across experimental phases. At Baseline 1, grouped subjects spent significantly less time in the 10 mm perimeter than isolated subjects (Grouped: 22. 5% ± 0. 5%, Isolated: 38. 2% ± 3. 1%, *p* < 0. 001). At Baseline 2, the effect of the testing condition varied by subsequent treatment group assignment: for subjects assigned to the Control group, the difference between grouped and isolated subjects was not statistically significant (Grouped: 26. 0% ± 1. 2%, Isolated: 33. 5% ± 3. 9%, *p* = 0. 12, **Figure 4a**); however, for subjects assigned to the 1.6 µmol/L nicotine group, grouped subjects spent significantly less time in the perimeter than isolated subjects (Grouped: 24. 3% ± 2. 0%, Isolated: 38. 3% ± 4. 9%, *p* = 0. 02, **Figure 4b**). At the final CPP Test, untreated grouped controls spent significantly less time in the perimeter than isolated controls (Grouped: 19. 9% ± 0. 7%, Isolated: 34. 3% ± 4. 1%, *p* = 0. 007, **Figure 4a**).

**Figure 4.**
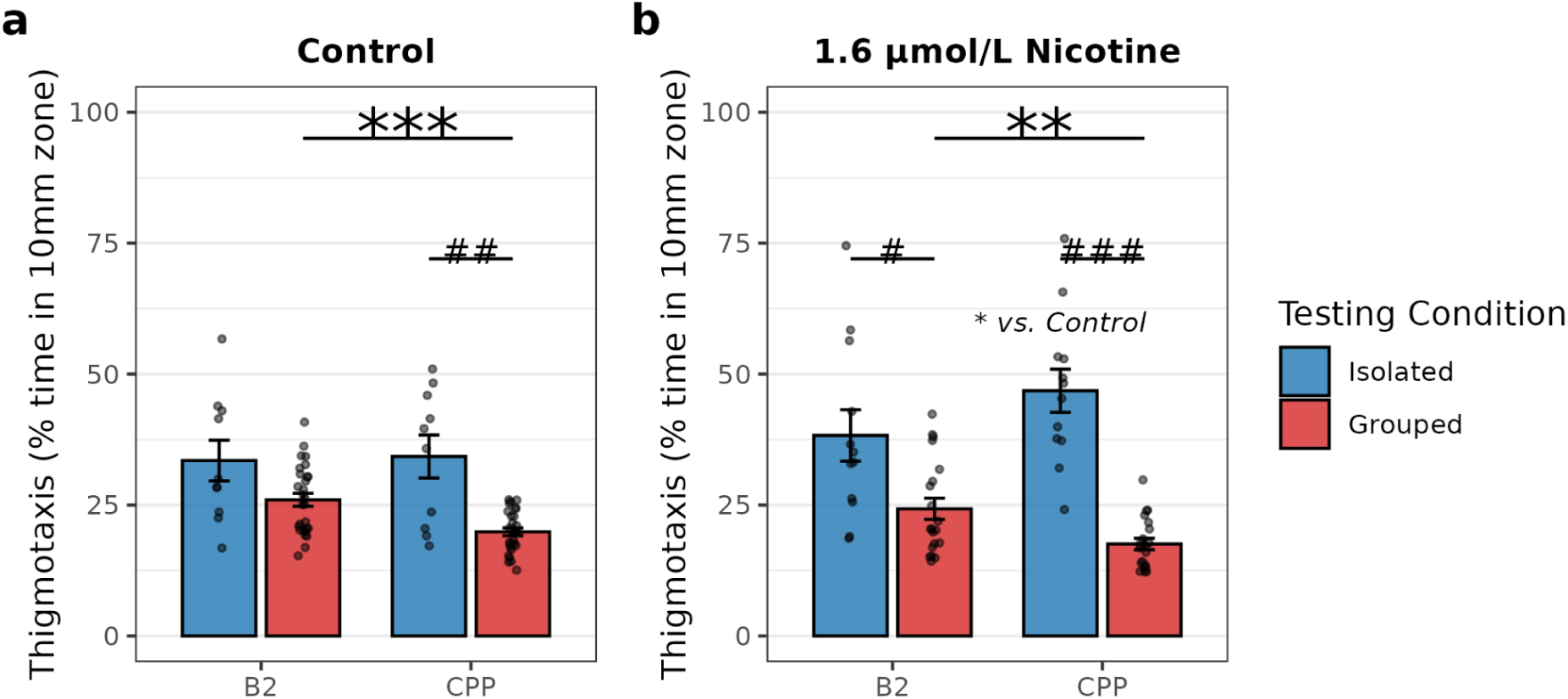
Thigmotaxis across experimental stages. Percentage of time spent in the 10 mm perimeter during Baseline 2 and the CPP Test for Control (left) and 1.6 µmol/L nicotine-treated (right) subjects. Grouped subjects showed significantly lower thigmotaxis than isolated subjects. Grouped subjects exhibited significant habituation (reduced thigmotaxis) over time regardless of treatment, whereas nicotine treatment significantly elevated thigmotaxis in isolated subjects at the final test compared to isolated controls. Data are presented as mean ± SEM. Significance markers: ** *p* < 0. 01, *** *p* < 0. 001 for stage contrasts (B2 vs CPP); # *p* < 0. 05, ## *p* < 0. 01, ### *p* < 0. 001 for testing condition contrasts (Grouped vs Isolated); * vs. Control indicates a significant difference (*p* < 0. 05) between the nicotine group and the corresponding Control group under isolated testing conditions at the CPP Test phase.

Longitudinal analysis across stages showed that the habituation trend in thigmotaxis was determined by the testing condition (social context) and was similar between the Control and nicotine-treated groups. Within the Control group, grouped subjects exhibited a significant decrease in thigmotaxis from Baseline 2 to the CPP Test (*p* < 0. 001), whereas isolated controls showed no significant change (*p* = 0. 79). Similarly, in the 1.6 µmol/L nicotine-treated group, grouped subjects displayed a significant reduction in thigmotaxis from Baseline 2 to the CPP Test (*p* = 0. 001), whereas isolated nicotine-treated subjects showed no significant change (*p* = 0. 18). Because of these contrasting longitudinal profiles between testing conditions, the Stage × Testing Condition interaction was statistically significant for both the Control group model (*p* = 0. 03) and the 1.6 µmol/L nicotine group model (*p* = 0. 02).

Although both treatment groups followed a similar longitudinal pattern, a significant Treatment × Testing Condition interaction was detected at the CPP Test phase (*p* = 0. 03). Simple effects analysis revealed that nicotine conditioning had divergent effects on thigmotaxis depending on the social context during testing. For grouped subjects, conditioning with the 1.6 µmol/L dose of nicotine resulted in a thigmotaxis level of 17. 6% ± 1. 1% (**Figure 4b**), which did not differ significantly from the 19. 9% ± 0. 7% observed in grouped controls (**Figure 4a**, *p* = 0. 32). In contrast, isolated subjects conditioned with the 1.6 µmol/L nicotine dose exhibited a thigmotaxis level of 46. 8% ± 4. 1% (**Figure 4b**), which was significantly elevated compared to the 34. 3% ± 4. 1% observed in isolated controls (**Figure 4a**, *p* = 0. 04). These findings indicate that while testing in a social group promotes habituation and reduces anxiety-like perimeter occupancy regardless of drug exposure, solitary testing represents a significant stressor that potentiates anxiety-like thigmotaxis following nicotine conditioning (**Figure 4**).

Across all cohorts and testing conditions, subjects exhibited no strong population-level innate bias toward either compartment (Isolated B1: 0.43 ± 0.03; Grouped B1: 0.47 ± 0.01) (**Figure 5**). We assessed the test-retest reliability of baseline preference across the two sessions. For the grouped condition, stability was evaluated at the group level by calculating the correlation between tank-averaged B1 and B2 scores. Grouped subjects demonstrated a high degree of behavioral stability, with a significant positive correlation between their tank-averaged baseline preferences (*r* = 0. 593, *p* = 0. 0059). In contrast, because isolated subjects were tested individually, their stability was evaluated at the individual level. Isolated subjects exhibited no significant correlation between their individual baseline scores across the two sessions (*r* = 0. 121, 95% CI: −0.32 to 0.52, *p* = 0. 590), indicating high intra-individual variability across days, although the small sample size (*N* = 22) limits the statistical power to rule out a moderate correlation (**Figure 6**).

**Figure 5.**
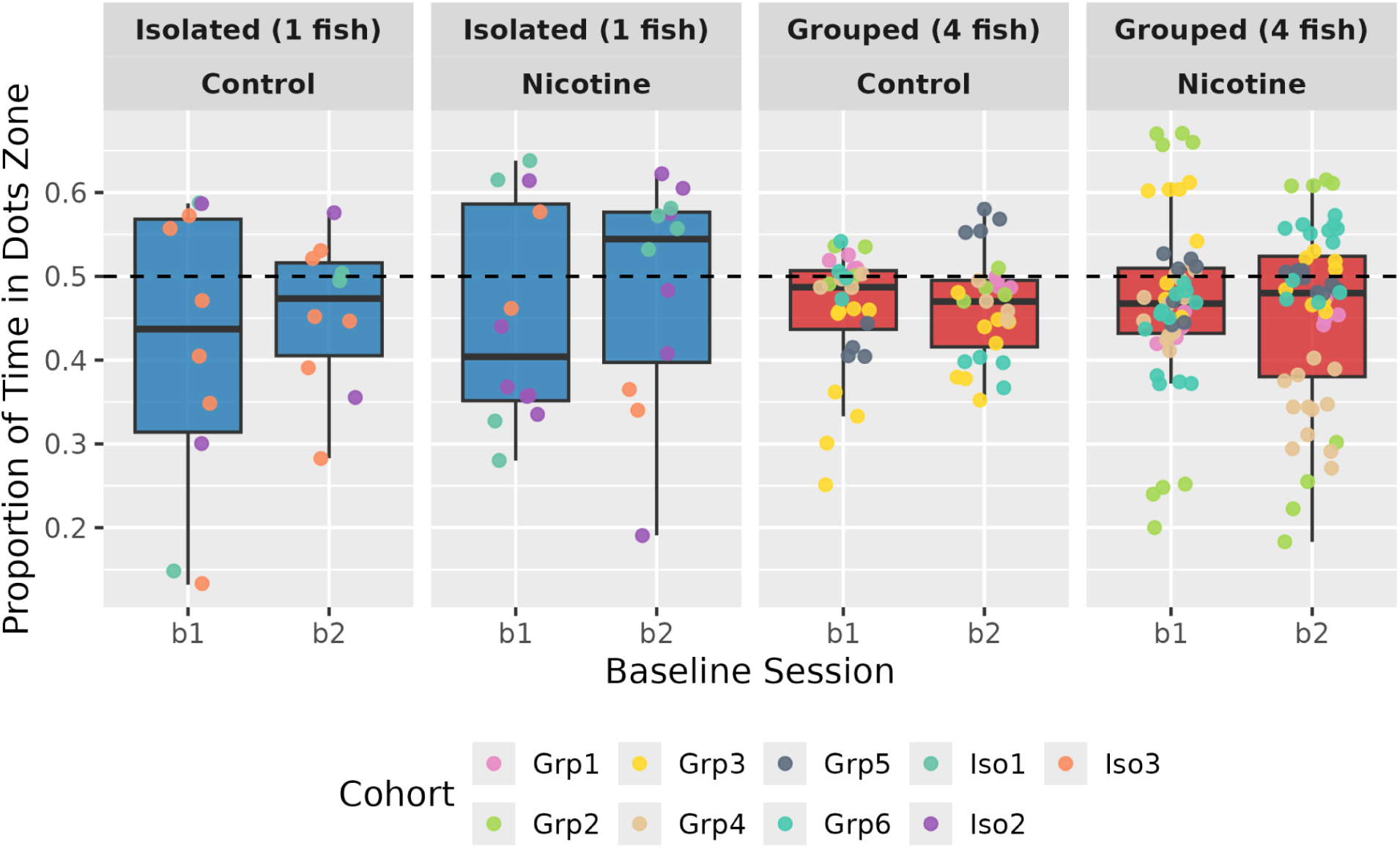
Baseline preference across cohorts. Distribution of the proportion of time spent in the dots-paired zone during the two baseline sessions (B1 and B2) across all experimental cohorts. No innate preference was observed for either the dotted or grid compartment.

**Figure 6.**
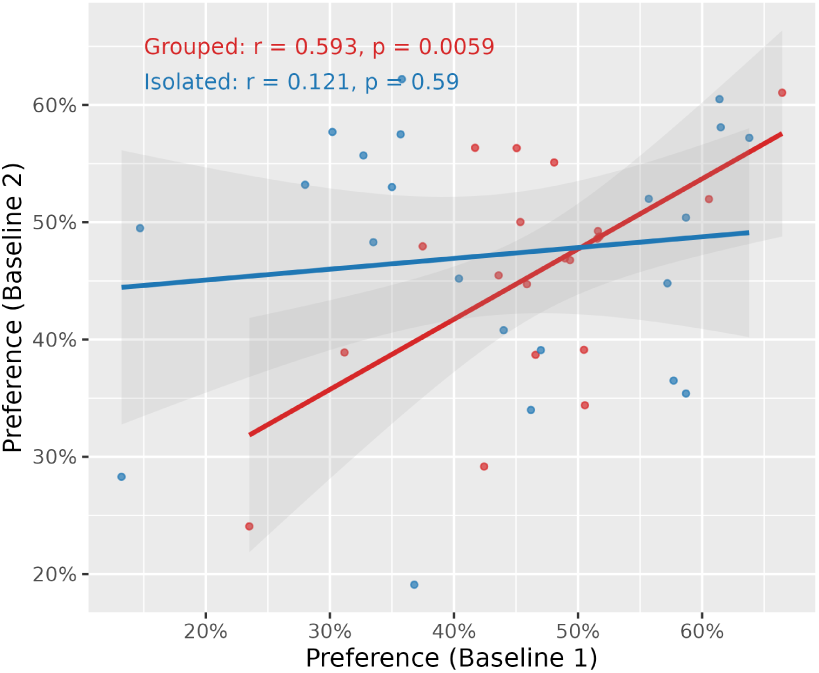
Test-retest reliability of baseline preference. Grouped subjects showed significant test-retest reliability in floor cue preference between Baseline 1 and Baseline 2 (*r* = 0. 593, *p* = 0. 0059), whereas isolated subjects showed no significant correlation (*r* = 0. 121, 95% CI: −0.32 to 0.52, *p* = 0. 590). The group-mean preference was used for group-tested fish as individual identities were not tracked. Shaded bands represent the 95% confidence interval.

To investigate the social dynamics driving this stability, we analyzed the variance structure of baseline preferences using a random-intercept LMM for the grouped fish. This analysis partitioned the total baseline preference variance into between-tank variance (τ^2^ = 0. 00862, Std. Dev. = 0. 093) and within-tank variance (σ^2^ = 0. 00036, Std. Dev. = 0. 019). The resulting Intraclass Correlation Coefficient (ICC) was 96. 0%, demonstrating that 96. 0% of the variance in baseline preference was between-tank variance, while only 4. 0% was individual variance within the tank. This confirms that grouped fish acted as highly cohesive, synchronized units within each tank.

To determine whether group testing restricted individual choices or homogenized behavior, we compared the overall population-level dispersion between testing conditions. The standard deviation of the tank averages in the grouped condition (*SD* = 0. 093, *N* = 20 tanks) was comparable to the population standard deviation of isolated individual fish (*SD* = 0. 114, *N* = 22 fish). Statistical tests confirmed there was no significant difference in overall dispersion between the two populations (Levene’s test, *p* = 0. 22; F-test, *p* = 0. 21). This demonstrates a “high-cohesion, high-diversity” structure, where grouped fish exhibit strong behavioral cohesion at the tank level while fully preserving the natural behavioral diversity of the species across the population (**Figure 7**).

**Figure 7.**
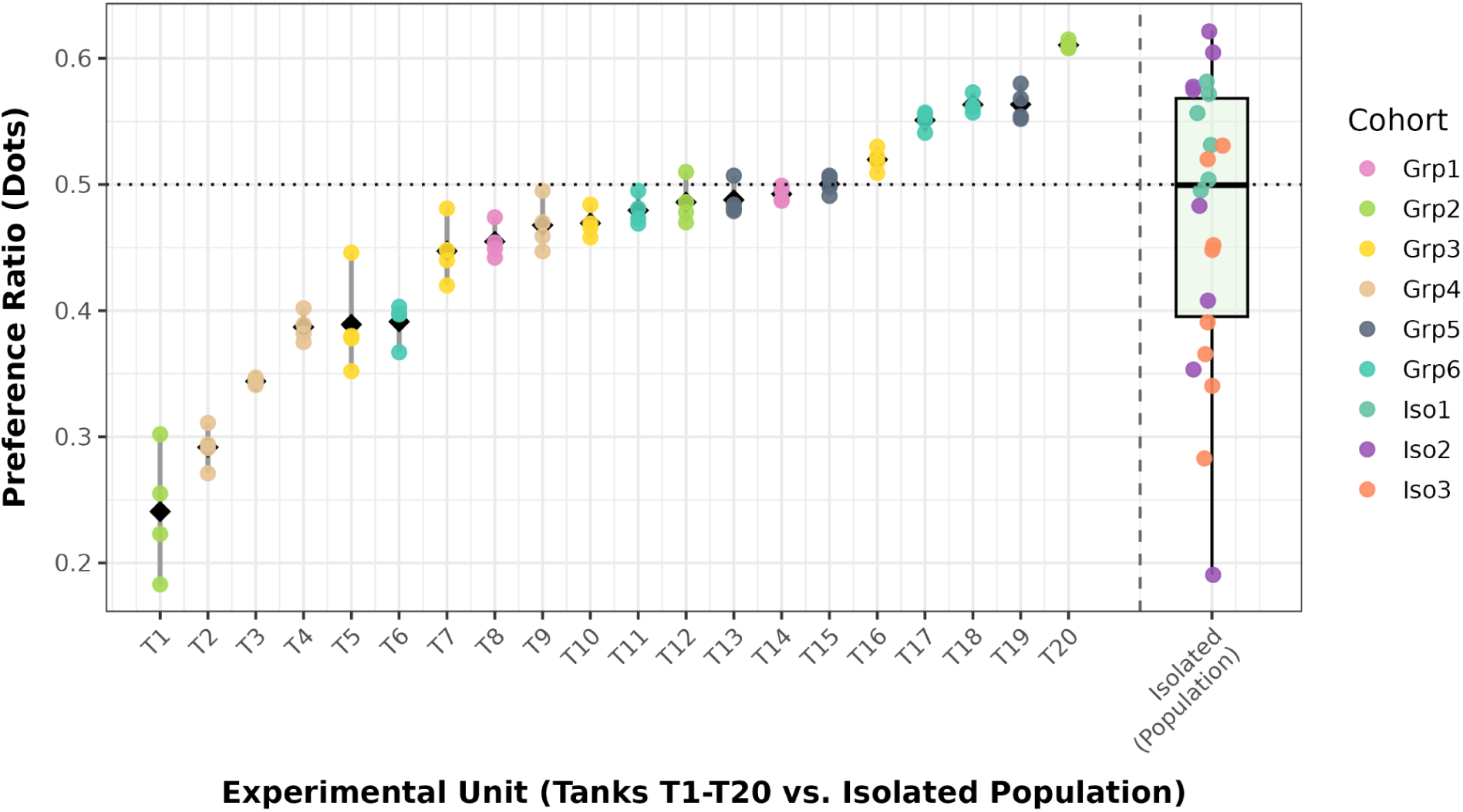
Preference variability and population dispersion. Comparison of inter-individual variability (standard deviation of the preference ratio) between grouped (intra-tank) and isolated (within-group) contexts across treatments. Dispersion in the grouped condition (tank averages) did not differ significantly from the population dispersion of isolated individual subjects.

### Conditioned Place Preference

To determine whether nicotine could induce a conditioned place preference, we analyzed the shift in preference toward the initially non-preferred (stimulus-paired) compartment following conditioning. Because individual identity cannot be maintained in the grouped condition across sessions, we utilized a LMM with Tank ID as a random intercept to account for shared environmental variance. In the grouped condition (4 fish/tank), the LMM revealed that the 1.6 µmol/L nicotine dose induced a statistically significant increase in preference for the paired compartment relative to the control group (*Estimate* = 0. 131, *SE* = 0. 051, *t*_15.79_ = 2. 568, *p* = 0. 0208). The 5.0 µmol/L dose also produced a significant preference shift (*Estimate* = 0. 129, *SE* = 0. 060, *t*_15.79_ = 2. 156, *p* = 0. 0468), whereas the lowest dose (0.5 µmol/L) showed a marginal trend (*p* = 0. 0630).

As a secondary analysis, we tested whether each treatment group’s CPP score differed from zero (i.e., whether the group shifted from its own baseline). In the grouped condition, all three nicotine doses produced CPP scores that were significantly greater than zero (0.5 µmol/L: *t*_15.9_ = 3. 11, *p* = 0. 007; 1.6 µmol/L: *t*_15.8_ = 3. 86, *p* = 0. 001; 5.0 µmol/L: *t*_15.8_ = 2. 96, *p* = 0. 009), whereas the grouped control group did not shift from baseline (*t*_15.8_ = 0. 58, *p* = 0. 569) (**Figure 8**).

**Figure 8.**
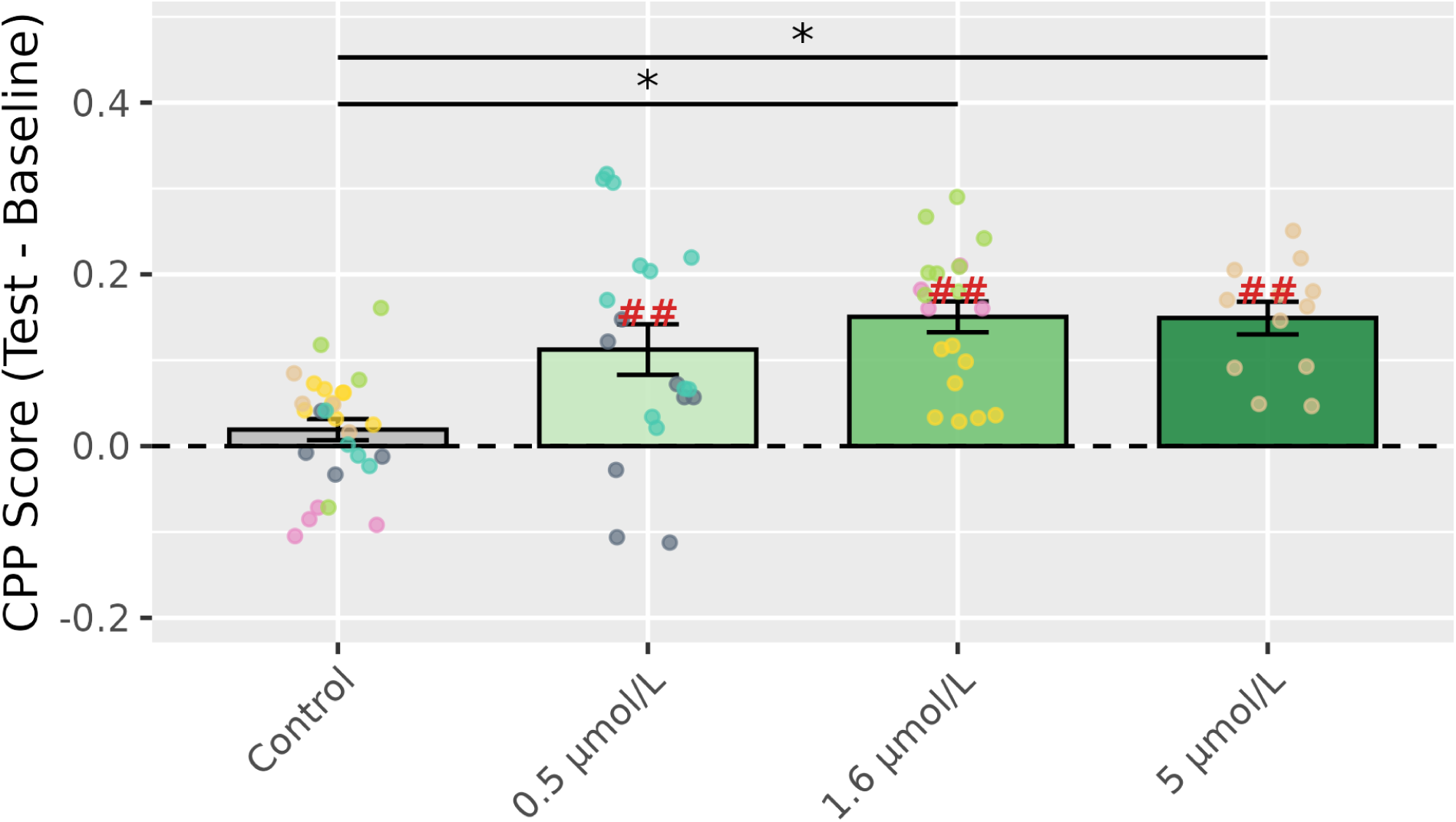
Nicotine-induced place preference in grouped subjects. Shift in preference toward the nicotine-paired compartment following conditioning in grouped subjects (4 fish/tank). Black brackets with asterisks denote significant differences from the control group (between-group comparison, * *p* < 0. 05). Red hash marks above individual bars denote CPP scores that differ significantly from zero (within-group comparison, ## *p* < 0. 01). Data are presented as mean ± SEM.

Conversely, in the isolated condition (1 fish/tank), a standard linear model did not detect a significant place preference shift in the surviving isolated cohort receiving the 1.6 µmol/L nicotine dose compared to controls (*Estimate* = 0. 044, 95% CI: −0.030 to 0.118, *SE* = 0. 038, *t*_20_ = 1. 176, *p* = 0. 253). However, the within-group analysis revealed that both the isolated control and nicotine groups shifted significantly from their own baselines (control: *t*_9_ = 3. 39, *p* = 0. 008; 1.6 µmol/L: *t*_11_ = 6. 97, *p* < 0. 001). The positive shift in the isolated control group (mean = 0.110) indicates a systematic baseline drift toward the initially non-preferred compartment, likely reflecting habituation to the apparatus over repeated sessions rather than a drug effect. Because this drift was present in both treatment groups, it was controlled for in the between-group comparison (nicotine vs. control), which remains the primary analysis. The smaller sample size and attrition in the isolated condition further limit the statistical power to draw definitive negative conclusions (**Figure 9**).

**Figure 9.**
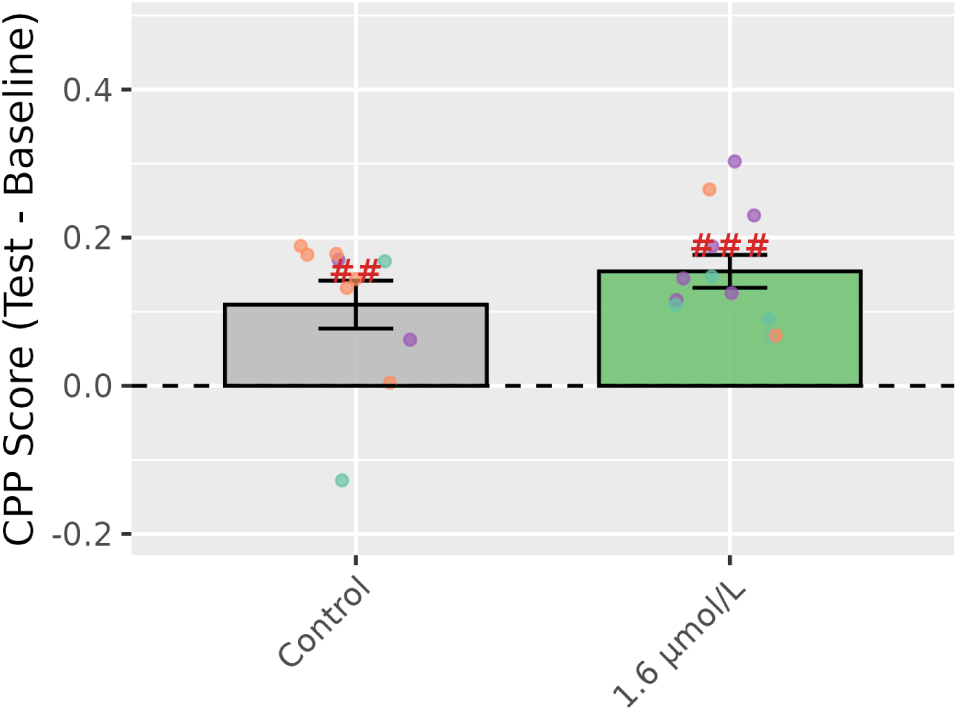
Nicotine-induced place preference in isolated subjects. Shift in preference following conditioning in isolated subjects (1 fish/tank) tested at 1.6 µmol/L nicotine. No significant difference was observed between the nicotine and control groups (between-group comparison). Red hash marks above individual bars denote CPP scores that differ significantly from zero (within-group comparison, ## *p* < 0. 01, ### *p* < 0. 001). The significant shift in the control group suggests a systematic baseline drift in isolated subjects. Data are presented as mean ± SEM.

### Locomotor Activity and Trajectory

Locomotor activity, measured as total distance traveled and mean swimming speed, was evaluated to ensure that preference shifts were not confounded by general locomotor changes (such as drug-induced hyperactivity or sedation), and to assess locomotor differences related to the testing social context. To appropriately account for shared environmental variance in grouped subjects and differences in variability between testing conditions, longitudinal locomotor metrics were analyzed using LMM with Tank ID as a random intercept and heterogeneous residual variances modeled across testing conditions in the statistical models.

At Baseline 2, cross-sectional analysis revealed that controls tested in groups were significantly more active than controls tested individually in total distance (*p* = 0. 01, Figure 10a) and mean swimming speed (*p* = 0. 01, Figure 10c), suggesting that isolated testing may induce a comparatively hypoactive state in naturally shoaling zebrafish. In the cohort designated for 1.6 µmol/L nicotine treatment, this baseline difference between testing conditions was not significant (*p* > 0. 05; Speed: *p* > 0. 05, Figure 10b, 10d).

**Figure 10.**
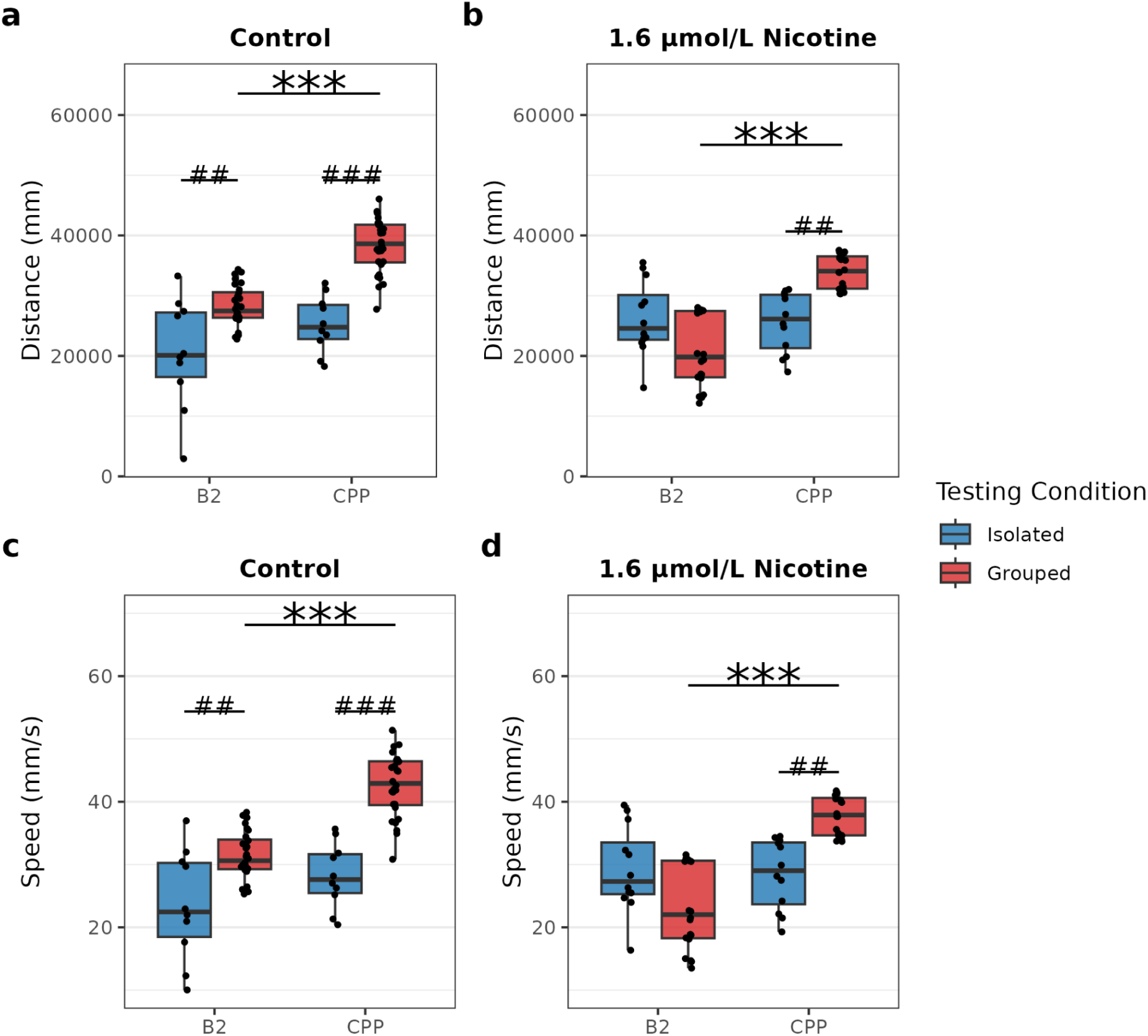
Locomotor activity across experimental stages. Total distance traveled (top) and average swimming speed (bottom) during Baseline 2 and the CPP Test for control and 1.6 µmol/L nicotine-treated subjects. Grouped subjects showed a highly significant increase in activity during the test, whereas isolated subjects remained hypoactive.

Longitudinal analysis across stages revealed significant shifts in activity following conditioning, which were dependent on social context. Within the Control groups, grouped fish exhibited an increase from Baseline 2 to the CPP Test in both total distance (*p* < 0. 0001, Figure 10a) and swimming speed (*p* < 0. 0001, Figure 10b). In contrast, isolated controls did not exhibit a significant change in activity (Distance: *p* = 0. 13, Figure 10a; Speed: *p* = 0. 13, Figure 10c).

An identical pattern emerged in the 1.6 µmol/L nicotine-treated groups. Grouped subjects displayed highly significant increases in both total distance (*p* < 0. 0001, Figure 10b) and swimming speed (*p* < 0. 0001, Figure 10d) during the CPP test. In contrast, isolated fish receiving the exact same 1.6 µmol/L nicotine dose exhibited no change in their locomotor activity (Distance: *p* = 0. 69, Figure 10b; Speed: *p* = 0. 68, Figure 10d). Together, these results suggest that both handling-induced and nicotine-associated locomotor changes are modulated by social context.

## Discussion

This study demonstrates that social context determines behavioral stability, experimental feasibility, and the expression of conditioned reward in juvenile zebrafish. Grouped subjects tested in cohorts of four exhibited a robust, dose-dependent place preference for the nicotine-paired compartment, with the most pronounced preference expressed at the 1.6 µmol/L dose. In contrast, isolated subjects did not exhibit a significant place preference at the same dose as controls. This lack of detectable preference in isolated subjects was accompanied by a 38.9% experimental attrition rate and pronounced locomotor suppression. These results establish that the solitary testing environment serves as a confounding stressor that disrupts cognitive and behavioral readouts in juvenile zebrafish.

The high attrition and elevated thigmotaxis observed in isolated subjects indicate that solitary testing represents a significant anxiogenic stimulus. Isolated control subjects remained near the perimeter of the tank and displayed frequent freezing behaviors, whereas the presence of conspecifics during grouped testing reduced these anxiety-like responses and permitted normal exploratory behavior. These behavioral patterns align with neural mapping studies showing that a 48-hour social isolation period in juvenile zebrafish alters activity in hypothalamic and preoptic regions, inducing social aversion and freezing behavior that can be rescued by the anxiolytic drug buspirone (Tunbak et al. 2020). These developmental findings are also consistent with observations in adult zebrafish, where short-term individual housing can evoke anxiety-like responses, although behavioral and endocrine measures often decouple (Marchetto et al. 2021; Onarheim et al. 2022). Thus, the absence of detectable place preference in isolated subjects may represent a masking effect of isolation-induced anxiety and locomotor inhibition on preference retrieval, rather than a failure of primary nicotine reinforcement. Alternatively, isolation stress may directly interfere with the neural substrates of reinforcement learning; for instance, stress-induced cortisol elevation can blunt phasic dopamine responses in reward circuits, which could disrupt the acquisition of the conditioned association. Further research is required to distinguish between behavioral masking and stress-induced disruption of reward signaling. However, the smaller sample size and high attrition rate in the isolated condition inherently limit the statistical power to draw definitive negative conclusions regarding reward expression under solitary testing.

The secondary within-group analysis, which tested whether each treatment group’s CPP score differed from zero, provided additional insight. In grouped subjects, all three nicotine doses produced CPP scores significantly above zero, whereas the grouped control group showed no shift from baseline. This pattern confirms that the preference shifts in grouped nicotine-treated subjects reflected genuine conditioned learning rather than non-specific drift. In contrast, isolated subjects in both the control and nicotine groups shifted significantly from their own baselines. The positive shift in isolated controls (mean CPP score = 0.110, *p* = 0. 008) indicates a systematic baseline drift toward the initially non-preferred compartment across repeated sessions, possibly reflecting habituation to the less-preferred cue or increased exploratory sampling of the novel compartment. This drift inflated the apparent CPP score in all isolated subjects equally. Because the between-group comparison (nicotine vs. control) controlled for this shared drift, the non-significant between-group result (*p* = 0. 253) indicates that nicotine did not produce a preference shift beyond what was already present in isolated controls. The absence of comparable drift in grouped controls suggests that the presence of conspecifics stabilizes spatial preferences across sessions, consistent with the higher test-retest reliability observed in the grouped condition.

A key finding of this study is the characterization of the social dynamics governing the grouped testing environment. Variance partitioning of baseline preference in grouped fish revealed that 96.0% of the total variance was between-tank variance, while only 4.0% was individual variance within each tank. This partitioning confirms that grouped fish acted as cohesive, synchronized units during the sessions. However, comparison of the overall population-level dispersion showed that the standard deviation of tank averages in the grouped condition was comparable to the population standard deviation of isolated individuals. These findings demonstrate a high-cohesion, high-diversity social structure. Grouped testing stabilizes behavior at the tank level while fully preserving the natural, population-wide individual behavioral diversity of the species.

In this study, juvenile zebrafish expressed robust conditioned place preference at low-micromolar nicotine concentrations (0.5, 1.6, and 5.0 µmol/L), with the most pronounced reward expression at 1.6 µmol/L. These effective doses are substantially lower than those typically used in adult zebrafish place conditioning paradigms, which range from 5 to 50 µmol/L (or 15 to 50 mg/L) (Kedikian et al. 2013; Viscarra et al. 2020; Pisera-Fuster et al. 2021). The heightened sensitivity of younger zebrafish to low nicotine doses aligns with developmental findings in larval assays, where the rewarding effects operate within a narrow concentration window (0.63 to 6.3 µmol/L), and higher doses trigger sensory irritation or avoidance (Schneider et al. 2023). Notably, while the 1.6 µmol/L dose elicited the strongest preference, a significant effect was also observed at 5.0 µmol/L, and a positive trend was visible at 0.5 µmol/L. This dose-response pattern suggests that late juvenile zebrafish possess a broader window for nicotine reinforcement than larvae, where the rewarding effects are extremely narrow and easily masked by sensory irritation or toxicity at higher concentrations. Furthermore, while previous adult studies have utilized genetically variable commercial pools or local strains (Faillace et al. 2018; Viscarra et al. 2020), this study utilized the standard AB laboratory strain. The use of a genetically well-characterized, standardized line like the AB strain is essential for reducing genetic background noise and ensuring reproducibility across laboratories. Critically, because the vast majority of transgenic and knockout zebrafish lines are maintained on the AB genetic background, establishing the feasibility of this assay in this specific strain directly facilitates downstream molecular, transgenic, and genetic investigations into the neurobiology of addiction.

Several limitations of this study should be noted. First, although we hypothesized that conspecific presence serves as a social buffer to mitigate testing stress, the underlying physiological mechanisms (such as whole-body cortisol levels or neural stress markers) were not directly measured. Second, individual identities were not tracked across independent daily sessions in the grouped condition because individual fish could not be visually distinguished in the video recordings, requiring the use of tank-mean preferences for longitudinal calculations. Third, our evaluations were restricted to the standard AB laboratory strain, and the generalizability of these findings to other zebrafish strains remains to be established. Fourth, only a single nicotine dose (1.6 µ mol/L) was evaluated in the isolated condition due to cohort size and resource constraints. Consequently, we cannot determine whether the lack of statistically detectable place preference in isolated subjects is dose-dependent or represents a universal blockade of reward expression under solitary testing conditions.

In conclusion, these results suggest that accounting for the social environment is important when evaluating reward and locomotor behaviors in social organisms. Testing juvenile zebrafish in groups of four provides a reliable, low-attrition, and ethologically relevant paradigm. This design stabilizes spatial exploration and is associated with more reliable expression of conditioned place preference. This grouped testing format represents a methodological improvement for developmental behavioral pharmacology and substance abuse research in zebrafish.

## Declarations

### Ethics Approval

All animal experimental procedures were approved by the Institutional Animal Care and Use Committee (IACUC) of the University of Tennessee Health Science Center and were conducted in accordance with the National Institutes of Health Guide for the Care and Use of Laboratory Animals.

### Consent to Participate

Not applicable.

### Consent for Publication

Not applicable.

### Availability of Data and Materials

The datasets generated and analyzed during the current study are available from the corresponding author upon reasonable request.

### Code Availability

Software code controlling recording of video and 3D printing of test chamber and visual cues are available at https://github.com/miraclezero/FishCPP.

### Authors’ Contributions

JH, TV, and HC contributed to the study design, protocol development, and experimental setup. JH performed the behavioral experiments. JH and HC analyzed the video tracking data and coordinate trajectories. All authors contributed to writing the manuscript, revised it for important intellectual content, and approved the final version.

